# CYB561 supports the neuroendocrine phenotype in castration-resistant prostate cancer

**DOI:** 10.1101/2024.02.29.582710

**Authors:** Romie Angelo G. Azur, Kevin Christian V. Olarte, Weand S. Ybañez, Alessandria Maeve M. Ocampo, Pia D. Bagamasbad

## Abstract

Castration-resistant prostate cancer (CRPC) is associated with resistance to androgen deprivation therapy, and an increase in the population of neuroendocrine (NE) differentiated cells. It is hypothesized that NE differentiated cells secrete neuropeptides that support androgen-independent tumor growth and induce aggressiveness of adjacent proliferating tumor cells through a paracrine mechanism. The cytochrome b561 (*CYB561*) gene, which codes for a secretory vesicle transmembrane protein, is constitutively expressed in NE cells and highly expressed in CRPC. CYB561 is involved in the α-amidation-dependent activation of neuropeptides, and contributes to regulating iron metabolism which is often dysregulated in cancer. These findings led us to hypothesize that CYB561 may be a key player in the NE differentiation process that drives the progression and maintenance of the highly aggressive NE phenotype in CRPC. In our study, we found that *CYB561* expression is upregulated in metastatic and NE prostate cancer (NEPC) tumors and cell lines compared to normal prostate epithelia, and that its expression is independent of androgen regulation. Knockdown of *CYB561* in androgen-deprived LNCaP cells dampened NE differentiation potential and transdifferentiation-induced increase in iron levels. In NEPC PC-3 cells, depletion of CYB561 reduced the secretion of growth-promoting factors, lowered intracellular ferrous iron concentration, and mitigated the highly aggressive nature of these cells in complementary assays for cancer hallmarks. These findings demonstrate the role of CYB561 in facilitating transdifferentiation and maintenance of NE phenotype in CRPC through its involvement in neuropeptide biosynthesis and iron metabolism pathways.

## Introduction

Primary prostate cancer (PCa) development is predominantly driven by androgens acting through its cognate binding partner, the androgen receptor (AR) (1), and consequently led to the use of androgen-deprivation therapy (ADT) as the standard care regimen for PCa (2). While ADT is initially effective in alleviating disease burden, patients ultimately develop resistance to therapy and the tumor recurs as a more aggressive disease referred to as castration-resistant PCa (CRPC) (3). A small percentage of advanced CRPC cases undergo transdifferentiation and develop neuroendocrine (NE) characteristics (4). Known as treatment-induced NE PCa (t-NEPC), this AR-independent CRPC subtype is characterized by pervasive metastasis, indolent therapy response, and poor prognosis (4). The incidence rate of t-NEPC tumors has steadily increased with the use of more potent ADT drugs such as enzalutamide (Enz) or abiraterone (5) which underscores the importance of deciphering the molecular underpinnings behind t-NEPC. One defining characteristic of t-NEPC tumors is the ability to secrete a wide spectrum of neuropeptides that promote cancer progression through a paracrine or autocrine manner (6), implicating neuropeptide signaling in acquiring resistance to ADT and driving the development of more aggressive tumors.

The integral membrane protein cytochrome b561 (CYB561) is a key component in neuropeptide synthesis and activation. In NE vesicles, CYB561 reduces monodehydroascorbate into ascorbic acid (ASC) which is then used by peptidylglycine α-amidating monooxygenase (PAM) for the α-amidation step required for neuropeptide activation (7). CYB561 also influences intracellular iron metabolism (8), another dysregulated pathway in various cancer types (9). CYB561 exhibits ferrireductase activity by reducing endocytosed ferric (Fe^3+^) iron to the more catalytic ferrous (Fe^2+^) form that constitutes the cytosolic labile iron pool (LIP) (10). The elevated expression of *CYB561* in CRPC relative to metastatic PCa (11, 12), along with its dual role in neuropeptide signaling and iron metabolism, led us to explore a possible contextual relationship between CYB561, and transformation and maintenance of NEPC. Notably, the possible involvement of CYB561 in relation to CRPC development and NEPC has not been functionally established.

In this study, we conducted *in silico* analysis to determine the pattern of *CYB561* expression across different prostate tumor subtypes, and characterized the basal and androgen-dependent regulation of *CYB561* expression in *in vitro* cell models representative of normal prostate epithelium, androgen-dependent metastatic disease, hormone-indolent CRPC, and NEPC. We found that *CYB561* expression is higher in castration-resistant and NE prostate tumors and cell lines, and is independent of androgen regulation in the AR-expressing metastatic LNCaP cell line. Knockdown of *CYB561* in LNCaP and induction of NE transdifferentiation through chronic steroid-starvation enhanced sensitivity to Enz, altered the expression of genes involved in neuropeptide synthesis and NE differentiation (NED), and decreased Fe^2+^concentration in parallel with changes in the expression of iron-responsive genes (IRGs). Knocking down *CYB561* in the NEPC cell model PC-3 decreased the expression of NED markers and secretion of growth-promoting factors, lowered the intracellular Fe^2+^concentration, affected IRG expression, and reduced overall aggressive cancer cell behavior based on several cancer hallmarks. To our knowledge, this is the first study to establish the functional involvement of CYB561 in CRPC development and NEPC.

## Materials and Methods

### Cell culture

PNT1A, RWPE-1, LNCaP, 22Rv1, and PC-3 cells were used to model PCa progression. PNT1A is a normal post-pubertal human prostate epithelial cell line immortalized with SV40 (13). RWPE-1 is a non-neoplastic adult human prostatic epithelial cell immortalized with human papillomavirus 18 (14). LNCaP is a model of androgen-responsive, metastatic prostatic adenocarcinoma that expresses AR and prostate specific antigen (PSA) (15). LNCaP cells also express the T878A mutated form of AR which can be transactivated by estrogens and other ligands, and may confer resistance to first line ADT drugs (16). The 22Rv1 cell line, a model of the progression of androgen-dependent prostate adenocarcinoma (PRAD) to NEPC, is a xenograft-derivative of CWR22 PRAD cells after serial propagation in castrated mice (17). Like LNCaP, 22Rv1 cells also express AR and PSA but has weak response to dihydrotestosterone (DHT) and androgen analogues. PC-3 cells have a NE background, closely resemble cells present in small cell NE carcinoma (18), do not express AR and PSA, exhibit highly aggressive behavior, and have been widely used to represent castration-resistant tumors (19). PNT1A (Cat. No. 95012614) and LNCaP (Cat. No. 89110211) cells were purchased from the European Collection of Authenticated Cell Cultures, while 22Rv1 cells were obtained from the American Type Culture Collection (Cat. No. CRL-2505). RWPE-1 and PC-3 cells were obtained from Dr. Diane Robins of the University of Michigan Medical School. The PC-3 cell line was authenticated by Macrogen (Korea) using short tandem repeat (STR) profiling (Powerplex 21 System, Promega). All cell lines tested negative for mycoplasma contamination using the Microsart AMP Mycoplasma Kit (Sartorius).

The cell lines PNT1A, LNCaP, 22Rv1, and PC-3 were maintained in RPMI 1640 medium (Gibco, 31800-022) supplemented with 10% fetal bovine serum (FBS; Gibco, 10500064), and 1X penicillin-streptomycin (Gibco, 10378-016). RWPE-1 cells were maintained in Keratinocyte Serum Free Medium (KSFM) (Gibco, 17005-042) supplemented with bovine pituitary extract and epidermal growth factor (Gibco, 37000-015). All cell lines were cultured under a humidified atmosphere of 5% CO_2_ at 37°C.

### Hormone treatment

For gene expression analyses, cells were seeded in 12-well plates (2.5 × 10^5^ cells/well) coated with poly-D-lysine (Gibco, A3890401). Hormone treatments were done upon reaching 70-80% confluency. For experiments requiring androgen starvation, LNCaP and 22Rv1 cells were grown in RPMI 1640 supplemented with 10% charcoal steroid-stripped (CSS) FBS for at least 48 hr (acute CSS; aCSS) prior to hormone treatment. Cells were then treated with 10 nM DHT (0.0058% methanol; Sigma, D-073-1ML) or 10 nM DHT plus 10 μM Enz (0.006% DMSO; Sigma, PHB00235) for 24 hr prior to RNA extraction. The control treatment received equivalent concentrations of vehicle solution.

To model the transdifferentiation of PRAD to NEPC, shRNA-scrambled and sh*CYB561*-transduced LNCaP cells were grown in RPMI 1640 with 1% CSS for at least 14 days as previously described (20). To complement this approach and simulate NEPC progression through long-term androgen starvation, we also maintained parental LNCaP cells in 10% CSS growth media for 10 months (chronic steroid starvation-cCSS). Induction of t-NEPC was confirmed by increased expression of NE gene marker, synaptophysin (*SYP*) (S1A Fig) (21) and formation of neurite-like extensions (S1B Fig) (22).

### *In silico* analyses of neuropeptide receptor gene expression in prostate cancer

We analyzed publicly available gene expression datasets to determine the expression pattern of *CYB561* across different tumor types. Using the ONCOMINE database (23), we examined the Bittner Multi-cancer clinical microarray data set (GSE2109) from the International Genomics Consortium Project for Oncology (www.intgen.org) for *CYB561* mRNA expression across different clinical tumor types. We also analyzed publicly available RNA-seq data on PRAD from the Cancer Genome Atlas (TCGA) Research Network (www.cancer.gov/tcga) (24), and RNA-seq and microarray copy number data on CRPC and NEPC clinical tumor samples (25). GEPIA (26) and cBioPortal (27) were used to generate overexpression or amplification plots for *CYB561, PAM,* and representative neuropeptide receptors in the different PCa subtypes.

### RNA extraction, reverse transcription and quantitative PCR

Total RNA was extracted from cell lines using the TRIzol reagent (Invitrogen, 15596-018) following the manufacturer’s protocol. Synthesis of cDNA and gene expression analysis were performed as described previously (28). Primers used to quantify mRNA levels were designed to span an exon-exon boundary (Table 1).

**Table 1.**
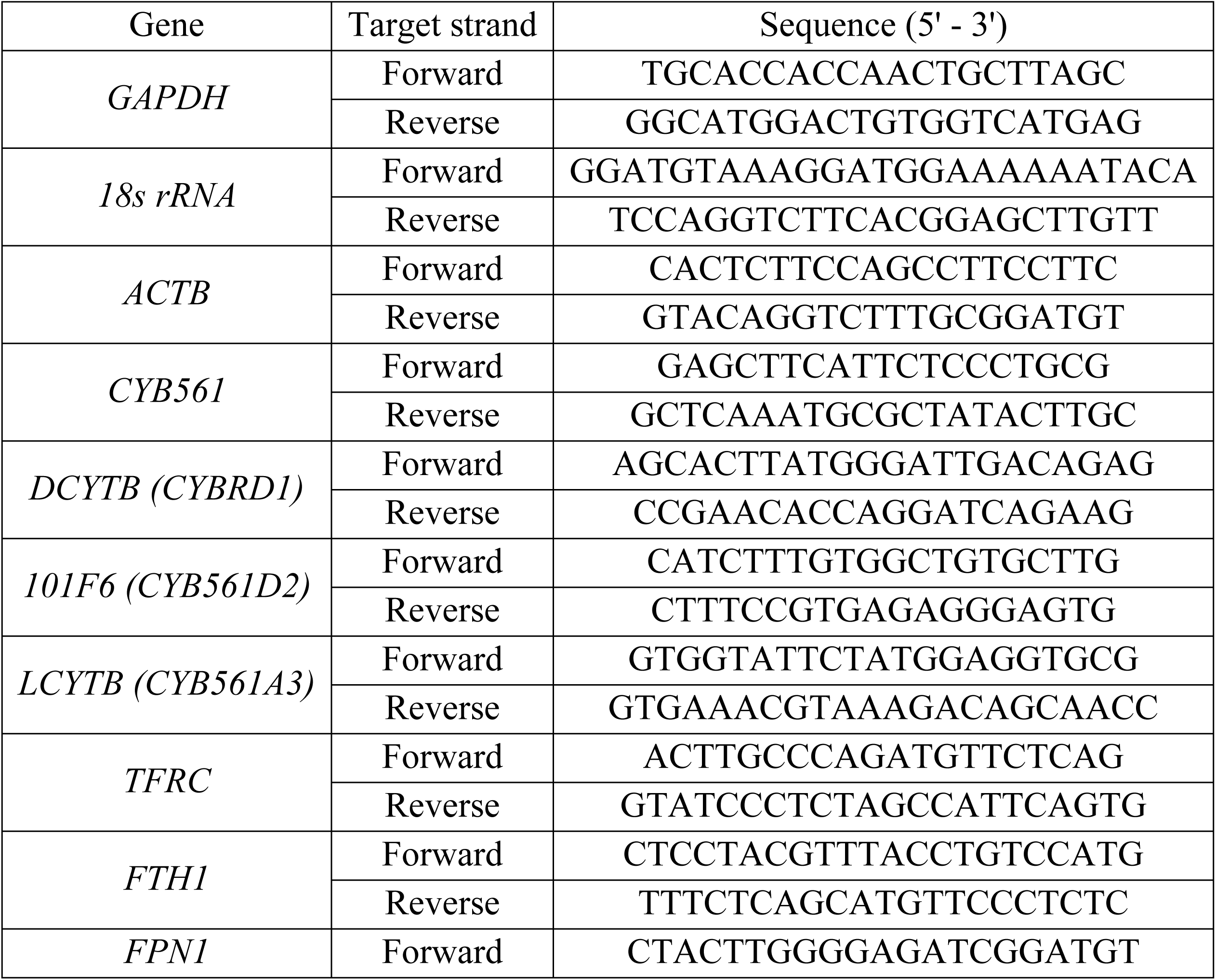
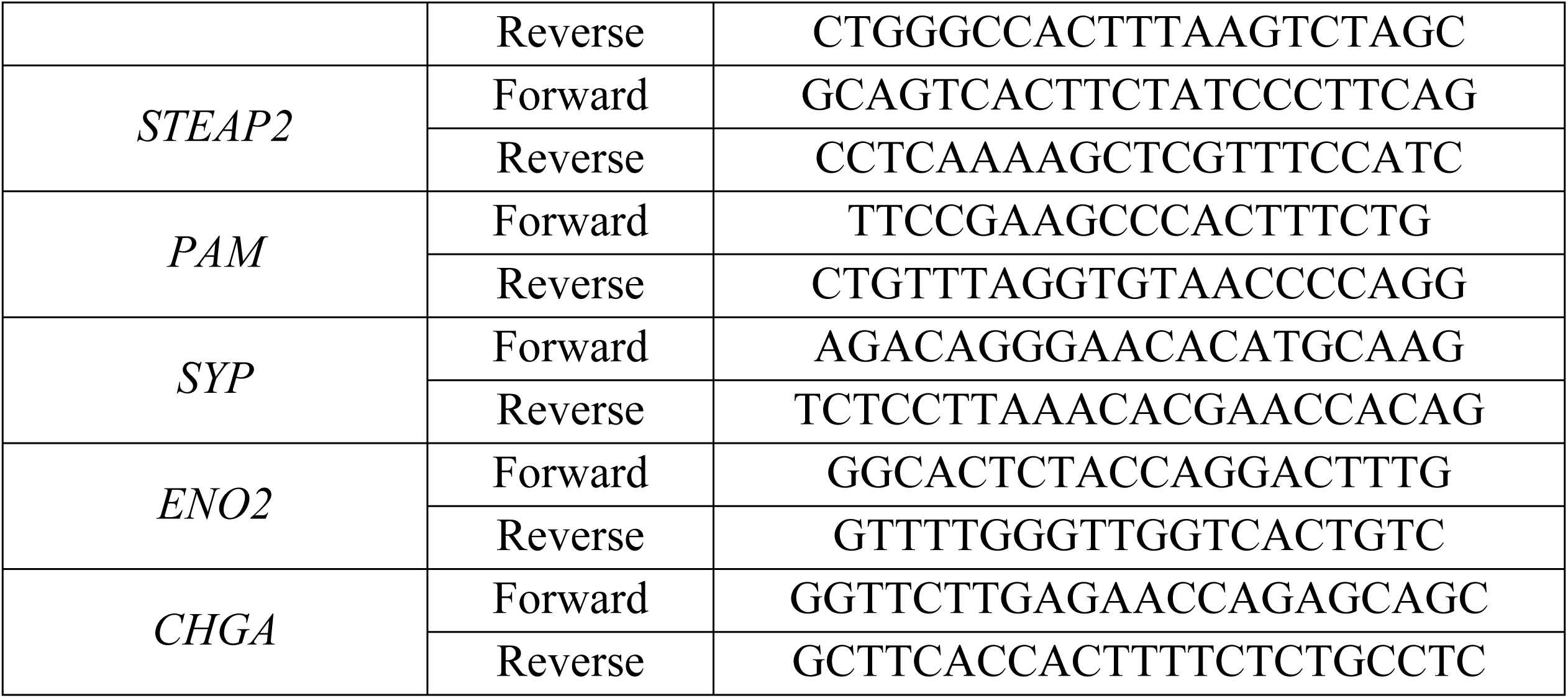
Primers used for RT-qPCR.

### Lentiviral-mediated knockdown of *CYB561*

The *CYB561* shRNA oligonucleotides (sh*CYB561*-1, TRCN0000439880; sh*CYB561*-3, TRCN0000064575; Genetic Perturbation Platform shRNA library, Broad Institute; S1 Table) were annealed and cloned into the pLKO.1 vector (RRID:Addgene_8453) (29) at the *Age*I and *Eco*RI cut sites.

Viral particles were packaged in HEK293T cells by transfecting the pLKO.1-sh*CYB561* constructs or scrambled shRNA pLKO.1 (RRID:Addgene_1864) (30) with viral packaging plasmids (pHCMVG, pRSV-Rev, pMDLg/pRRE), filtered and harvested as described previously (28). Transduced cells were selected using media with 2 µg/mL puromycin (Gibco, A11138-03). *CYB561* gene expression analysis by RT-qPCR was performed to verify successful knockdown.

### Trypan blue exclusion assay

PC-3 (5.0 × 10^4^ cells/well) transduced with scrambled shRNA control or sh*CYB561* were seeded in 6-well plates with complete media supplemented with 2 µg/mL puromycin. Cell cultures were allowed to proliferate for 5 days after which live cells were counted through trypan blue exclusion as previously described (28).

### CyQuant direct cell proliferation assay

PC-3 (1.0 × 10^3^ cells/well) and hormone-treated LNCaP (3.0 × 10^3^ cells/well) cells transduced with scrambled shRNA control or sh*CYB561* were seeded into 96-well clear-bottom black plates (Falcon, 353219) in complete or CSS media. Cell proliferation was monitored every 24 or 72 hr for 3-7 consecutive days using CyQuant Reagent (Invitrogen, C35011) by measuring bottom-read fluorescence at 520 nm with excitation wavelength set at 480 nm using EnSight Multimode Plate Reader (PerkinElmer). The same protocol was adapted to measure the effect of paracrine signaling on the proliferation rates of parental PNT1A (1.0 × 10^3^ cells/well), LNCaP (3.0 × 10^3^ cells/well), and 22rv1 (3.0 × 10^3^ cells/well) cells treated with conditioned media.

### Colony formation assay

PC-3 (500 cells/well) transduced with scrambled shRNA control or sh*CYB561* were seeded in 6-well plates and grown in complete media replenished every 3 days for a duration of 14 days prior to manual colony counting. Cells were fixed, stained, and analyzed as described previously (28). Colonies were manually counted using the Vision SX45 Stereomicroscope such that a cell colony is defined as a group of more than 50 cells.

### Wound healing assay

PC-3 (3 × 10^5^ cells/ml) transduced with scrambled shRNA control or sh*CYB561* were seeded in 24-well plates and grown in complete media supplemented with 2 µg/mL puromycin. Upon reaching 90-100% confluency, cells were grown in serum-free basal media for 24 hr. Cells were then pre-treated with 10 µg/mL Mitomycin C (Millipore, 475820) for 2 hr to stop cell division (31). A scratch wound was made using a P200 pipet tip and wound closure was monitored as described previously (28).

### Intracellular iron concentration measurement

LNCaP (2 × 10^6^ cells/dish) and PC-3 (2.5 × 10^5^ cells/well) cells transduced with scrambled shRNA control or sh*CYB561* were seeded in 12-well plates and grown in media supplemented with 2 µg/mL puromycin. Upon reaching 90-100% confluency, cells were washed with 1X PBS, and treated with 200 µM ferric ammonium citrate (FAC) (Sigma, F5879) for 4 hr. Intracellular iron concentration was measured using the Iron Assay Kit (Sigma, MAK025) following manufacturer’s protocol with the intracellular total iron and ferrous iron concentrations expressed as mmole/L or µmole/L.

### Paracrine signaling assay

PC-3 transduced with scrambled shRNA control or sh*CYB561* were grown in T-75 flasks with serum-free base RPMI 1640 media. Upon reaching 60-70% confluency, conditioned media from these cells were harvested for two consecutive days with daily fresh media replacement. Serum-free base RPMI 1640 media was also incubated in an empty cell culture flask to serve as a control. Collected conditioned and control media were centrifuged at 200 x*g* for 3 min at 4°C to pellet cell debris, and filter-sterilized using 0.22 µm Minisart® PES syringe filters (Sartorius, 16532-K). The filtered conditioned media from the transduced PC-3 were diluted 1:1 with fresh serum-free RPMI 1640 base medium. Control media were prepared by making a 1:1 dilution of the incubated control medium with fresh base medium (negative control) or 1:1 dilution of the incubated control medium with fresh base medium and supplementation with 2.5% FBS (positive control). Conditioned and control media were used as treatment media for PNT1A, LNCaP and 22Rv1 cells in downstream cell proliferation assays as previously described.

### Western Blot

Model cell lines were seeded at a density of 2.0 x 10^6^ cells per 100-mm culture dish and cultured until they reached 90% confluency. Total protein was extracted using RIPA buffer mixed with protease inhibitor cocktail (Millipore Merck, 539134) and quantified using Qubit® Protein Assay Kit (Thermo Fisher, Q33211). Proteins were separated by size by running 20 μg of the protein extract in 8% SDS-PAGE and transferred to Immuno-Blot PVDF Membrane (Bio-Rad, 1620174) using the Trans-Blot® SD Semi-Dry Transfer Cell (Bio-Rad) protocol. Non-specific interactions were blocked by incubating the membrane in TBS-T with 5% non-fat milk overnight at 4°C with constant agitation. To detect human CYB561 protein, an anti-CYB561 rabbit polyclonal antibody (Sigma-Aldrich, HPA014753) was used as the primary antibody and a goat polyclonal anti-rabbit IgG conjugated to horseradish peroxidase (Millipore, 12-348) for the secondary antibody. To visualize protein bands, the ECL™ Prime Western Blotting Reagent (Cytiva, RPN2236) was used and images captured using the iBright CL100 Imaging System (Invitrogen). A housekeeping protein was also detected using the same blot by mild stripping and re-probing the membrane with an anti-GAPDH rabbit polyclonal antibody (Sigma-Aldrich, G9545-200).

### Statistical Analysis

Results are presented as the mean ± SEM. Data for gene expression analysis (normalized to *18s rRNA*, *GAPDH,* or actin-beta (*ACTB*) transcript levels), wound healing assay (percentage relative to 0 hr), and cell proliferation (relative to signal at 0 hr) were log_10_ transformed before statistical analysis. The basal gene expression data were analyzed using one-way ANOVA followed by Tukey’s post hoc test. Gene expression data from hormone-treated LNCaP cells transduced with scrambled shRNA or sh*CYB561*, or grown in complete or CSS media were analyzed using two-way ANOVA to assess interaction of *CYB561*-knockdown or growth conditions and hormone treatment. Student’s unpaired *t*-test was then performed to analyze the effect of *CYB561*-knockdown or growth condition on hormone response. Results from transdifferentiation experiment were analyzed using two-way ANOVA to determine for main effects of transdifferentiation and *CYB561* knockdown, followed by Student’s unpaired *t*-test to determine the effect of transdifferentiation within an shRNA group and the effect of *CYB561* knockdown in cells maintained in similar growth conditions. Data from the hormone treatment and CSS gene expression analysis, and PC-3 *CYB561* knockdown lines experiments on colony formation, cell proliferation, manual cell count, wound healing, and iron measurement were analyzed using Student’s unpaired *t*-test. All statistical analyses were done using GraphPad Prism version 8.0 (GraphPad Software, La Jolla California USA, www.graphpad.com) and *P* < 0.05 was accepted as statistically significant.

## Results

### *CYB561* expression is elevated in castration-resistant prostate tumors and does not depend on androgen regulation

From our *in silico* analysis, we found that *CYB561* mRNA is upregulated in several cancer types, with the highest expression levels observed in prostate tumor samples (Fig 1A). *CYB561* is included in the top 3% of over-expressed genes in PCa tumors with an expression fold change of 2.064 compared to normal tissue equivalents. Analysis of publicly available RNA-seq and copy number data reflected a similar expression trend with higher *CYB561* mRNA levels in PRAD versus normal tissue samples (Fig 1B), and increasing *CYB561* amplification frequency as PCa progresses from PRAD to CRPC and eventually NEPC (Fig 1C). Expression of *PAM* (Fig 1D) and representative neuropeptide receptor genes (S2 Fig) also exhibited higher amplification frequencies in CRPC and NEPC compared to PRAD.

**Fig 1.**
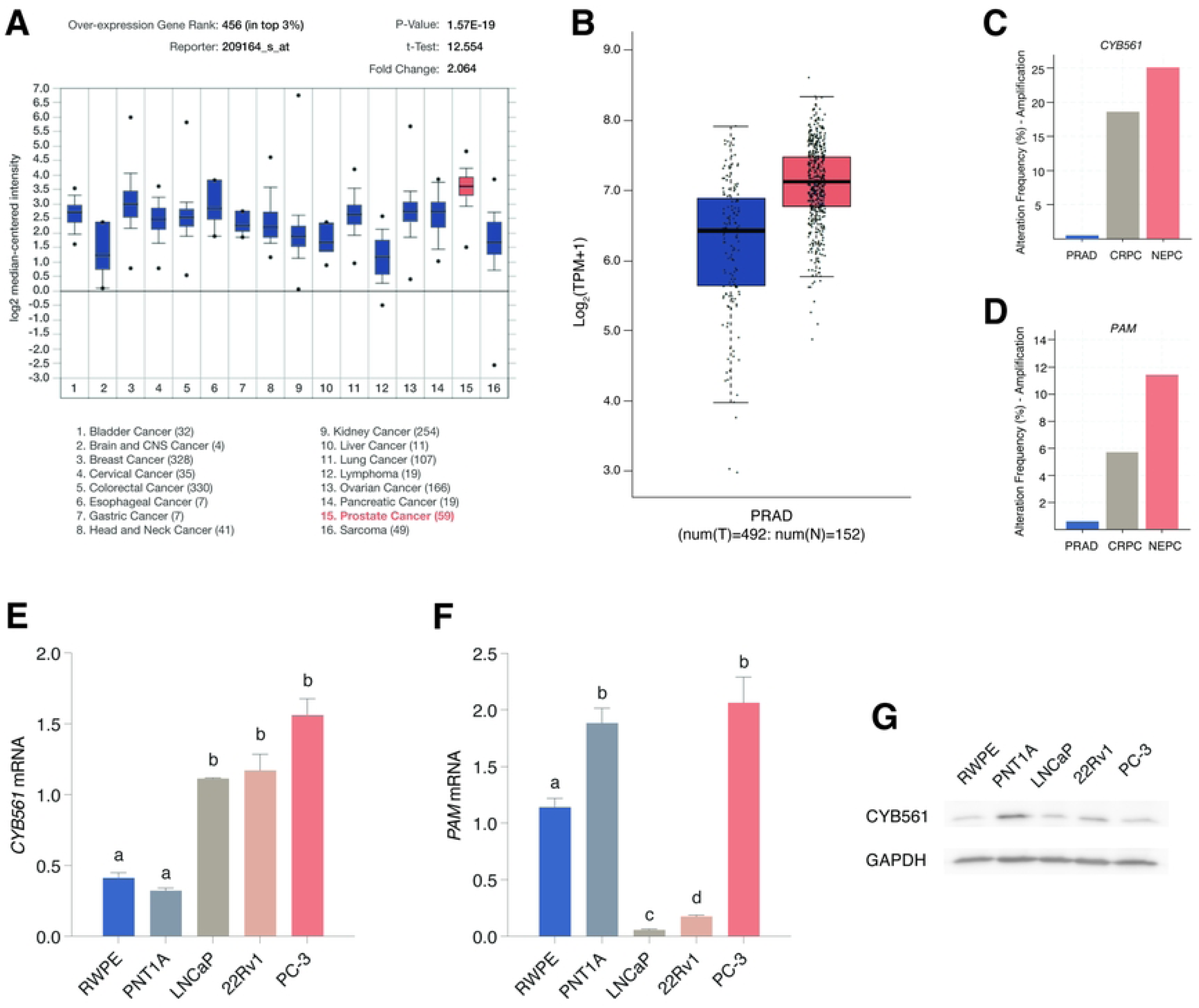
*CYB561* is highly expressed in several cancer tissue types and exhibits an increasing trend in expression as PCa progresses to CRPC and NEPC. (A) Publicly available microarray data were examined using the integrated data analysis and microarray repository, ONCOMINE (23). Based on the Bittner Multi-cancer microarray data set (GSE2109), *CYB561* expression was observed to be highest in prostate tumor samples and is included in the top 3% over-expressed genes in PCa compared to normal tissue equivalents. **(B)** Congruent analysis of RNA-seq data from the Genotype-Tissue Expression Project (GTEx) for normal prostate tissue (N) (32) and the Cancer Genome Atlas (TCGA) – prostate adenocarcinoma (PRAD; T) (33) using the online tool Gene Expression Profiling Interactive Analysis (GEPIA) (26) showed an increasing trend in *CYB561* transcript levels in PRAD samples compared to normal tissue equivalents. **(C-D)** The TCGA– PRAD (www.cancer.gov/tcga), and castration-resistant (CRPC) and neuroendocrine prostate cancer (NEPC) (25) RNA-seq and copy number data were analyzed using the online visualization tool cBioPortal (27, 34). Increasing (C) *CYB561* and (D) *PAM* expression correlates with disease severity and resistance to therapy, with highest amplification frequencies seen in CRPC and NEPC. **(E)** Baseline expression of *CYB561* mRNA is higher in the PCa cell lines LNCaP, 22RV1 and PC-3 compared to the normal prostate cell lines RWPE-1 and PNT1A. **(F)** Baseline expression of *PAM* mRNA is higher in the AR-negative cell lines RWPE-1, PNT1A, and PC-3 compared to LNCaP and 22rv1 cell lines. **(G)** Immunoblot for CYB561 across the five cell lines show that 22rv1 and PC-3 cells have higher CYB561 expression compared to LNCaP cells with PNT1A exhibiting the highest protein expression. Bars represent mean ± SEM with statistically significant differences determined through one-way ANOVA followed by Tukey’s post-hoc test (*P* < 0.05; means with the same letter are not statistically different).

To support our *in silico* findings, we measured the baseline expression levels of *CYB561* mRNA in prostate cell lines, and found that *CYB561* is elevated in metastatic LNCaP and 22Rv1 cells, and in the NEPC cell model PC-3 relative to normal prostate epithelial cell lines RWPE and PNT1A (Fig 1E). Among the *CYB561* homologs, duodenal *DCYTB,* tumor suppressor *101F6,* and lysosomal *LCYTB,* only *CYB561* was found to be consistently upregulated in PCa cell lines relative to normal epithelial cells (S3 Fig). We also observed high levels of *PAM* transcript in AR-negative cell lines RWPE-1, PNT1A, and PC-3 (Fig 1F). Lastly, western blot for CYB561 across the five cell lines showed higher CYB561 expression in 22rv1 and PC-3 cells compared to LNCaP cells, with the highest CYB561 protein expression in PNT1A suggesting that post-transcriptional regulation may play a role in modulating CYB561 protein expression (Fig 1G).

Since *CYB561* is upregulated in CRPC tumors and PCa cells, we determined whether its expression in LNCaP cells is regulated by androgens, and whether transcript levels are altered by acute and chronic androgen withdrawal. While we observed the expected increase in kallikrein-3 (*KLK3* or PSA) expression with DHT treatment that was abolished with Enz (S4A Fig), *CYB561* transcript levels did not change with DHT and Enz treatment (Fig 2A and S4B Fig). In contrast, the expression of *PAM* decreased with DHT treatment (Fig 2B) and increased upon acute androgen withdrawal (S4C Fig). On the other hand, both *CYB561* (Fig 2C) and *PAM* mRNA levels (Fig 2D) did not change in response to DHT or Enz treatment in 22Rv1 cells. Maintenance of LNCaP cells in chronic CSS conditions led to a loss of *KLK3* expression (S4D Fig), an increasing pattern of *CYB561* expression (S4E Fig), and a marked increase in *PAM* mRNA levels (S4F Fig). These data provide evidence that although *CYB561* expression is not dependent on androgen regulation, the neuropeptide synthesis pathway, through the induction of *PAM* expression, is activated upon androgen-withdrawal in LNCaP cells. However, this androgen-dependent regulation of *PAM* is no longer observed in 22Rv1 cells.

**Fig 2.**
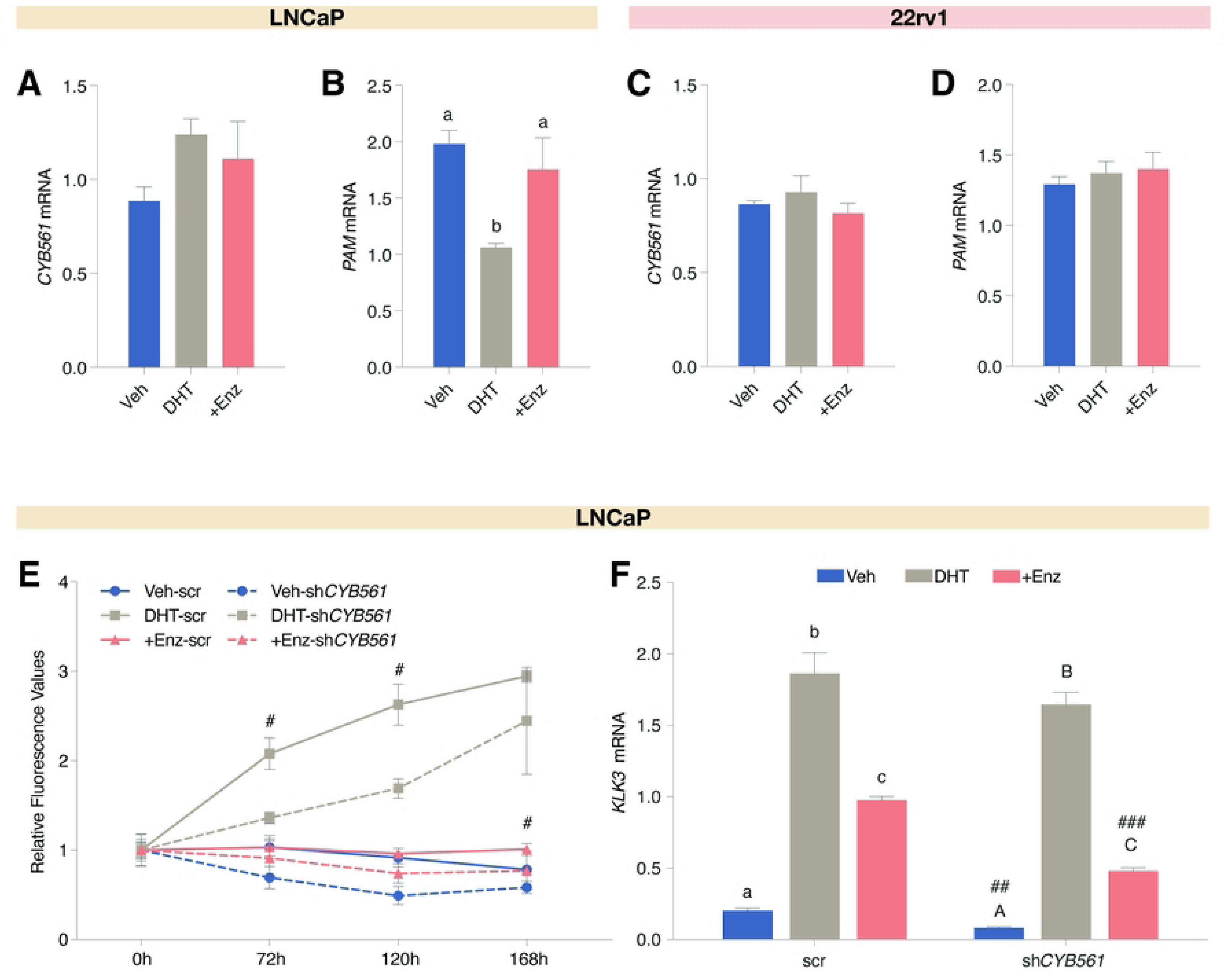
*CYB561* does not exhibit androgen-dependent gene regulation but contributes to androgen-sensitivity of LNCaP cells. **(A-B)** LNCaP and **(C-D)** 22Rv1 cells grown in hormone-starved media (aCSS) were treated with vehicle (Veh), 10 nM DHT, or 10 nM DHT plus 10 uM Enz for 24 hr before harvest and gene expression analysis by RT-qPCR. (A) *CYB561* mRNA expression was not affected by DHT and Enz treatment. (B) *PAM* mRNA expression decreased with DHT treatment and this effect was reversed in the presence of Enz. In 22Rv1, hormone treatment did not affect (C) *CYB561* and (D) *PAM* expression. **(E)** LNCaP cells transduced with the scrambled (scr) shRNA control and sh*CYB561* were treated with vehicle, 10 nM DHT, or 10 nM DHT plus 10 uM Enz, and cell proliferation was tracked for seven days. LNCaP cells transduced with the sh*CYB561* construct and treated with DHT and DHT plus Enz have lower cell proliferation rates compared to the corresponding scr shRNA control. **(F)** Expression of *KLK3* mRNA was induced in the scr and sh*CYB561* transduced cells upon DHT treatment and this effect was reduced with Enz. Notably, lower *KLK3* expression was observed for the Veh and Enz treated LNCaP-sh*CYB561* cells compared to the matched scr shRNA controls. Data points and bars represent mean ± SEM with statistically significant differences determined through one-way ANOVA followed by Tukey’s post-hoc test for effect of hormone treatment (*P* < 0.05; means with the same letter are not statistically different) and Student’s *t-*test for the effect of *CYB561* knockdown between the same hormone treatment (^#^*P* < 0.01, ^##^*P* < 0.001, ^###^*P* < 0.0001)

To determine whether CYB561 contributes to evading anti-PCa therapies, we knocked down *CYB561* expression in LNCaP cells, and measured cell viability and *KLK3* expression upon hormone treatment. Lentiviral-mediated transduction of sh*CYB561* in LNCaP cells effectively reduced *CYB561* mRNA expression by 74.24% relative to scrambled shRNA control (S5A Fig) which was also reflected at the protein level (S5B Fig), and remained unaltered with hormone treatment (S5C Fig). We found that *CYB561* knockdown reduced the proliferative effects of DHT (Fig 2E and S6A-B Fig) and enhanced the sensitivity of LNCaP cells to the anti-androgenic effects of Enz in terms of cell proliferation (Fig 2E and S6C Fig) and *KLK3* mRNA expression (Fig 2F).

### *CYB561* knockdown alters the expression of neuroendocrine markers and iron-responsive genes in transdifferentiated LNCaP cells

To determine the functional significance of *CYB561* upregulation in CRPC and t-NEPC development, *CYB561* knockdown LNCaP cells were maintained in serum-depleted growth conditions known to induce NE transdifferentiation (20). Similar to the effect of acute and chronic CSS growth conditions, transdifferentiation did not affect *CYB561* transcript levels in LNCaP cells (Fig 3A) but increased *PAM* mRNA expression which was further enhanced upon *CYB561* knockdown (Fig 3B). Expression of NED gene markers *SYP* (Fig 3C), neuron-specific enolase 2 (*ENO2*) (Fig 3D), and chromogranin A (*CHGA*) (S7 Fig) also increased upon transdifferentiation and these effects were dampened with *CYB561* knockdown.

**Fig 3.**
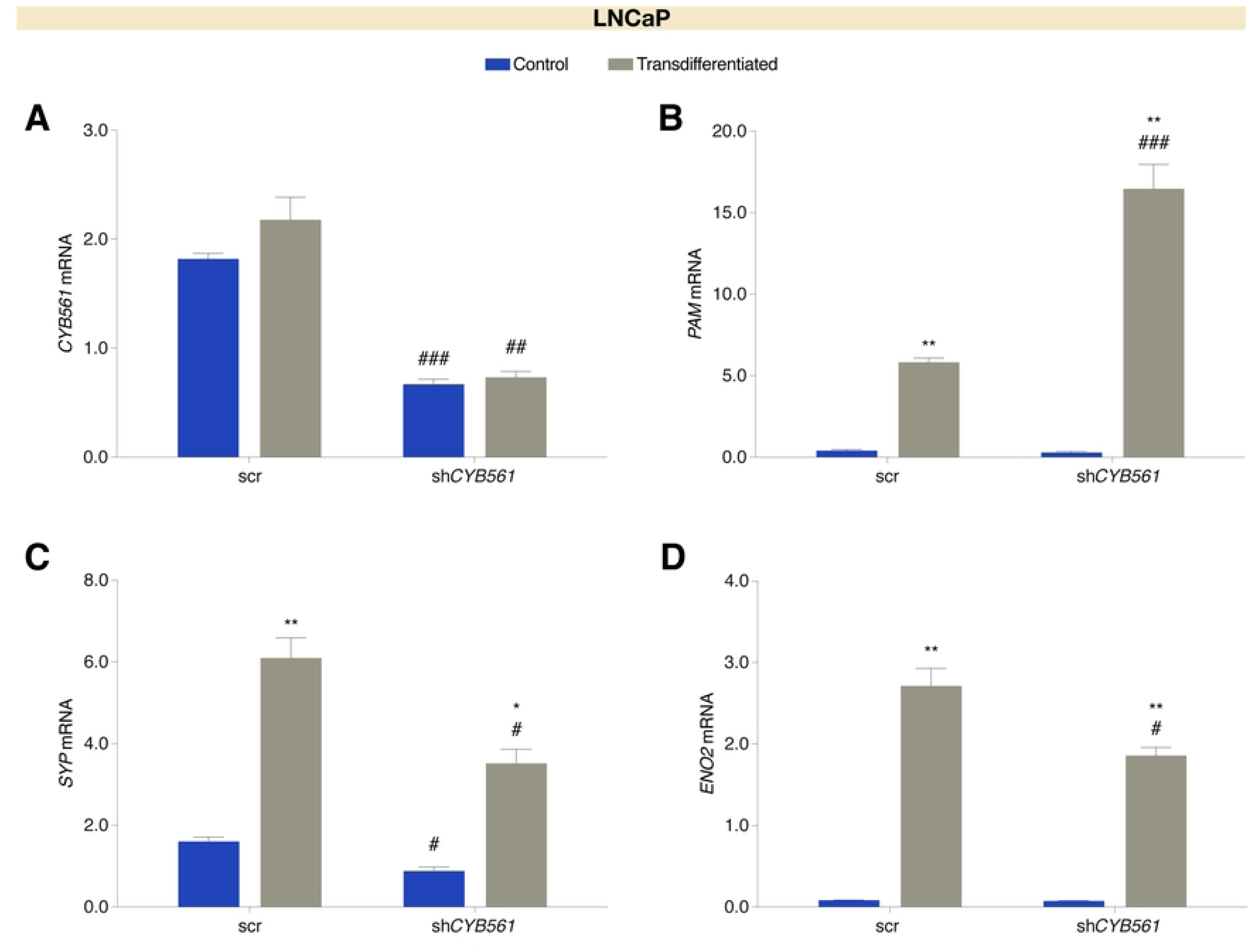
Expression of neuroendocrine differentiation gene markers is reduced upon *CYB561* knockdown in LNCaP cells. LNCaP cells transduced with the scrambled (scr) shRNA control and sh*CYB561* were grown and maintained in complete media (control) or transdifferentiation media for 14 days. **(A)** Transdifferentiation did not affect *CYB561* mRNA expression (two-way ANOVA; Treatment factor: *P* = 0.1079; Knockdown factor: *P* < 0.0001) while **(B)** *PAM* mRNA expression increased with transdifferentiation which was further enhanced with *CYB561* knockdown (two-way ANOVA; Treatment factor: *P* < 0.0001; Knockdown factor: *P* = 0.0791). The neuroendocrine differentiation gene markers **(C)** *SYP* (two-way ANOVA; Treatment factor: *P* < 0.0001; Knockdown factor: *P* < 0.0001) and **(D)** *ENO2* (two-way ANOVA; Treatment factor: *P* < 0.0001; Knockdown factor: *P* = 0.0016) mRNA expression increased with transdifferentiation and this effect was dampened with *CYB561* knockdown. Bars represent mean ± SEM with statistically significant differences determined through two-way ANOVA for main effects of treatment and *CYB561* knockdown and Student’s *t-*test for the individual effects of treatment within a shRNA type (\**P* < 0.001, \*\**P* < 0.0001) and *CYB561* knockdown between the same treatment (^#^*P* < 0.01, ^##^*P* < 0.001, ^###^*P* < 0.0001).

To establish the association between CYB561 and iron metabolism in the context of PCa progression, we first measured the baseline expression of IRGs, transferrin receptor protein 1 (*TFRC*), ferritin heavy chain (*FTH1*), and ferroportin (*SLC40A1* or *FPN1*) in prostate cell lines. *TFRC* codes for a transmembrane protein responsible for Fe^3+^ uptake while *FTH1* codes for the heavy chain of the main cellular Fe^3+^ storage protein ferritin (35) (Fig 4A). FPN1 functions as a basolateral exporter that facilitates the efflux of excess Fe^2+^ (35) (Fig 4A). We found that in comparison to normal prostate epithelial cells, LNCaP cells have higher basal levels of *TFRC* (S8A Fig) and *CYB561* mRNA (Fig 1E), similar basal levels of *FTH1* (S8B Fig) mRNA, and lower *FPN1* (S8C Fig) expression. Notably, LNCaP and 22Rv1 cells show a similar pattern of IRG expression except for the higher *FPN1* mRNA in 22rv1. PC-3 cells consistently showed highest basal expression of *TFRC* and *FTH* across all cell prostate cell lines.

**Fig 4.**
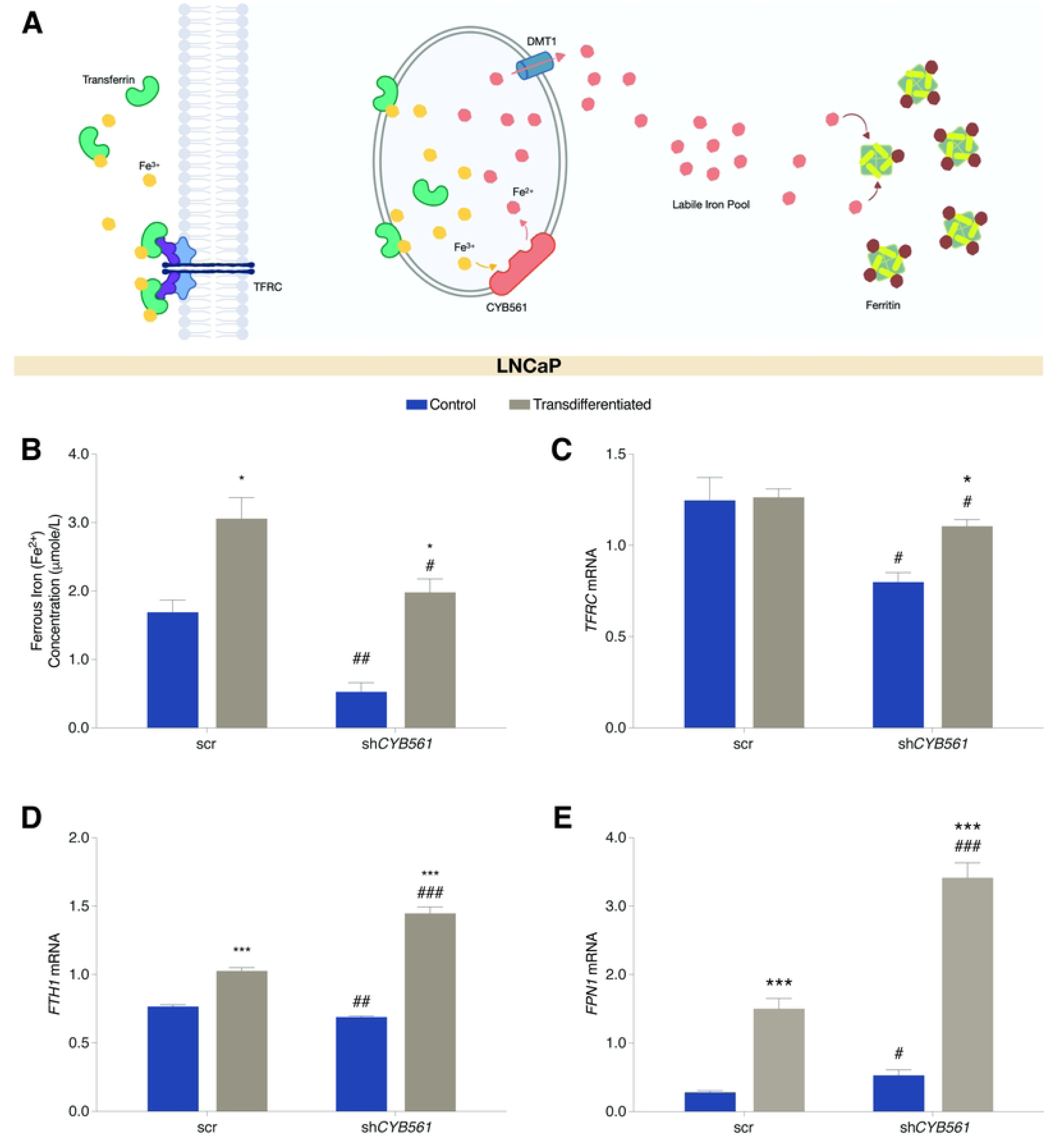
Transdifferentiation and *CYB561* knockdown altered the Fe^2+^ concentrations and expression of iron responsive genes (IRGs) in LNCaP cells. LNCaP cells transduced with the scrambled (scr) shRNA control and sh*CYB561* were grown and maintained in complete media (control) or transdifferentiation media for 14 days followed by Fe^2+^ concentration measurement. **(A)** CYB561 contributes to cellular iron homeostasis through its ferrireductase function to maintain vesicular redox states. This function is congruent with how other players regulate cellular iron, such as TFRC which facilitates extracellular Fe^3+^ uptake and ferritin (FTH) which controls Fe^3+^ storage within the cytosol. **(B)** Concentration of Fe^2+^ increased with transdifferentiation and this effect was dampened with *CYB561* knockdown (two-way ANOVA; Treatment factor: *P* = 0.0001; Knockdown factor: *P* = 0.0007). Expression analysis of IRGs showed that (C) *TFRC* mRNA expression decreased with *CYB561* knockdown in control and transdifferentiation media. However, *TFRC* expression increased in the LNCaP-sh*CYB561* cell line after transdifferentiation (two-way ANOVA; Treatment factor: *P* = 0.0220; Knockdown factor: *P* = 0.0013). **(D)** Expression of *FTH1* (two-way ANOVA; Treatment factor: *P* < 0.0001; Knockdown factor: *P* = 0.0002) and (E) *FPN1* (two-way ANOVA; Treatment factor: *P* < 0.0001; Knockdown factor: *P* < 0.0001) mRNA increased upon transdifferentiation of LNCaP-scr cells and this effect was enhanced with *CYB561* knockdown. Bars represent mean ± SEM with statistically significant differences determined through two-way ANOVA for main effects of media treatment and *CYB561* knockdown, and Student’s *t-*test for the individual effects of media treatment within an shRNA type (\**P* < 0.01, \*\**P* < 0.001, \*\*\**P* < 0.0001) and *CYB561* knockdown between the same media treatment (^#^*P* < 0.05, ^##^*P* < 0.01, ^###^*P* < 0.001).

To determine the role of CYB561 in iron metabolism in the context of t-NEPC development, we measured intracellular iron concentration and determined IRG expression pattern in transdifferentiated and sh*CYB561*-transduced LNCaP cells. Transdifferentiation increased Fe^2+^ (Fig 4B) and total iron concentrations (S8D Fig) and these effects were reduced with *CYB561* knockdown. Transdifferentiation did not affect *TFRC* in scrambled shRNA control cells but increased *TFRC* levels in *CYB561* knockdown cells (Fig 4C). Notably, *CYB561* knockdown also decreased the expression of *TFRC* relative to scrambled shRNA control cells. *FTH1* and *FPN1* mRNA expression in scrambled shRNA LNCaP cells were upregulated upon transdifferentiation and were further enhanced by *CYB561* knockdown (Figs 4D-E). Similar patterns of IRG expression were observed in the parental LNCaP cells maintained in chronic CSS condition (S8E-F Fig). We eliminated the possibility that the observed transdifferentiation-induced changes in IRG expression were driven by AR-dependent regulation since expression of IRGs did not appear to be androgen-responsive (S8G-H Fig). The expression level of another ferrireductase, *STEAP2,* was also measured to account for increases in Fe^2+^ concentration that cannot be attributed to changes in *CYB561* expression. We found that *STEAP2* mRNA increased upon transdifferentiation but this effect was not altered by *CYB561* knockdown (S8I Fig).

### CYB561 is involved in maintaining the neuroendocrine phenotype and iron metabolism of PC-3 cells

To provide a functional basis for the upregulation of *CYB561* in NEPC, we determined the effect of knocking down *CYB561* on the expression of *PAM* and NED markers in the NEPC model cell line PC-3. Lentiviral-mediated transduction of sh*CYB561* in PC-3 cells successfully reduced *CYB561* mRNA expression by 74.23% relative to scrambled shRNA control (S9A Fig) which was reflected at the level of protein expression (S9B Fig). *PAM* mRNA expression was not affected by *CYB561* knockdown (Fig 5A) while expression levels for NED markers *SYP* (Fig 5B), *ENO2* (Fig 5C), and *CHGA* (S9C Fig) decreased upon CYB561 depletion.

**Fig 5.**
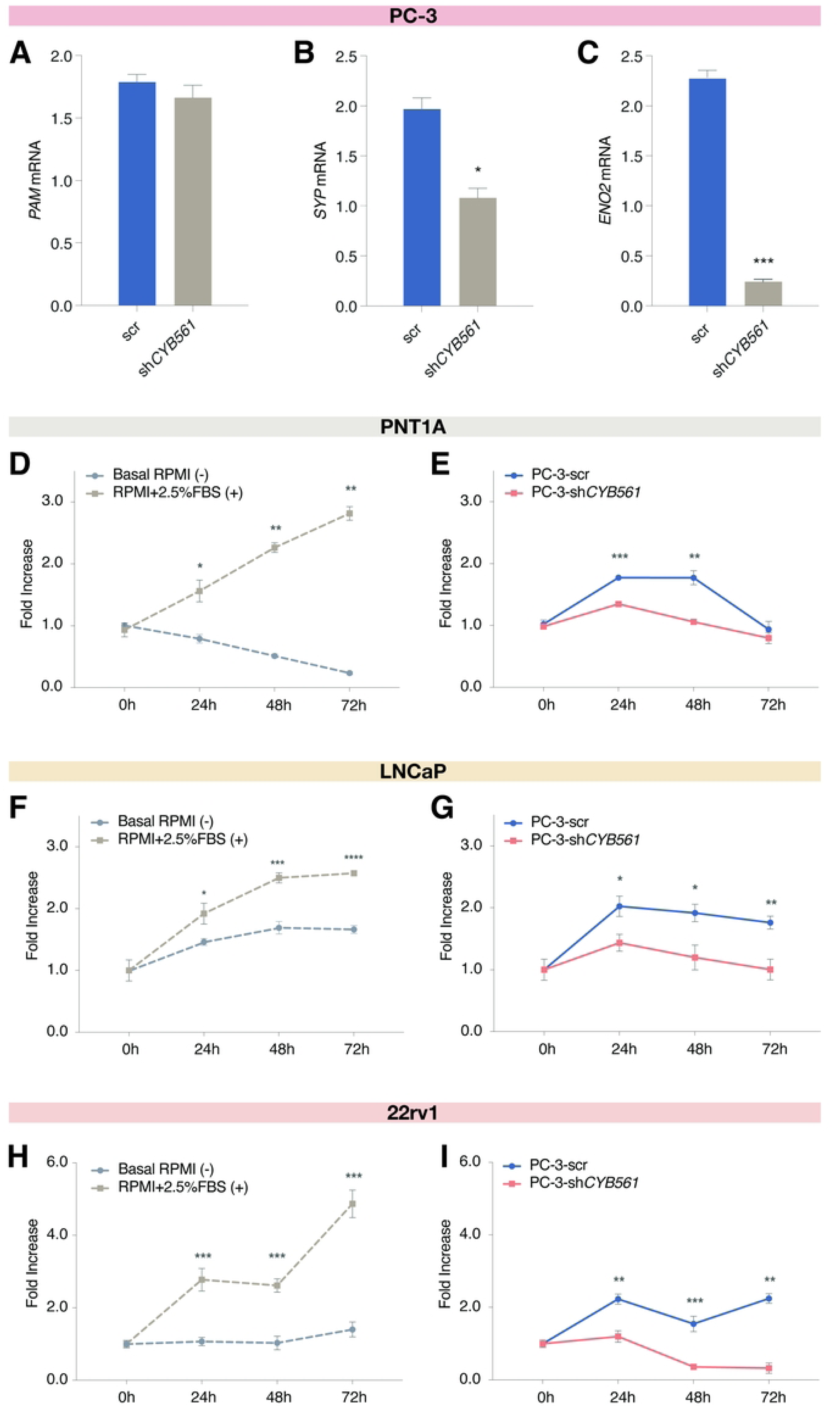
CYB561 contributes to the maintenance of the neuroendocrine phenotype of PC-3 cells. Knocking down *CYB561* expression had no effect on **(A)** *PAM* mRNA expression and downregulated the mRNA levels of the NED gene markers **(B)** *SYP* and **(C)** *ENO2*. **(D-E)** PNT1A, **(F-H)** LNCaP, and **(H-I)** 22rv1 cells were treated with conditioned media (CM) harvested from PC-3 cells transduced with scrambled (scr) shRNA control or sh*CYB561*, and cell survival was tracked for 2 days. Treatment of **(D)** PNT1A, **(F)** LNCaP, and **(H)** 22rv1 cells with the positive control media (base media supplemented with 2.5% FBS) was able to sustain the proliferation of PNT1A cells in contrast to the continuous decline in the proliferation of cells treated with the negative control media (base media only). Treatment of **(E)** PNT1A, **(G)** LNCaP, and **(I)** 22rv1 cells with the CM from the PC-3 scr shRNA control cells was able to support cell survival when compared to the CM from PC-3 cells transduced with sh*CYB561*. Bars and data points represent mean ± SEM with statistically significant differences determined by Student’s *t*-test (\**P* < 0.01, \*\**P* < 0.001, \*\*\**P* < 0.0001).

Since CYB561 is involved in paracrine signaling through its function in neuropeptide activation, we investigated the consequences of CYB561 knockdown on the secretion of paracrine factors that may support tumor growth. We used conditioned media derived from PC-3 scrambled shRNA control or sh*CYB561* cell lines as growth media for PNT1A prostate cells, LNCaP and 22Rv1 PCa cells. Treatment with conditioned media from PC-3 scrambled shRNA control and the positive control media were able to maintain cell survival and proliferation while treatment with PC-3 sh*CYB561*-derived conditioned media and negative control media led to a significant decline in the survival of cells across all three cell lines (Figs 5D-I). More importantly, cells treated with conditioned media from scrambled shRNA-transduced PC-3 cells exhibited higher survival rates than cells treated with conditioned media from sh*CYB561*-transduced PC-3 cells (Fig 5E, 5G, 5I).

To demonstrate the involvement of CYB561 in iron metabolism in the context of NEPC, we determined the effect of *CYB561* knockdown on intracellular iron concentrations and IRG expression in PC-3 cells. Knockdown of *CYB561* decreased Fe^2+^ levels (Fig 6A), led to a decreasing trend in total iron concentration (S9D Fig), upregulated *TFRC* expression (Fig 6B), and downregulated *FTH1* (Fig 6C) and *FPN1* (Fig 6D) expression.

**Fig 6.**
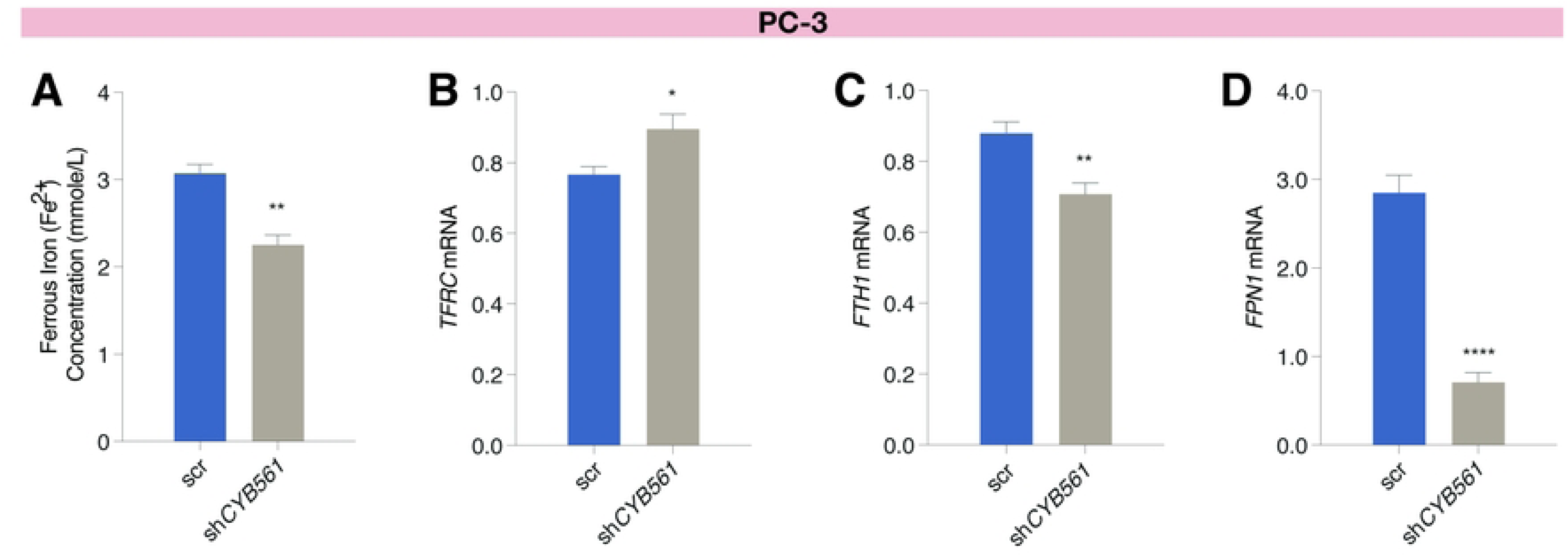
*CYB561* knockdown decreased intracellular Fe^2+^ iron concentration and altered the expression of iron responsive genes in PC-3 cells. Knockdown of *CYB561* in PC-3 cells **(A)** reduced intracellular Fe^2+^ concentration, **(B)** increased *TFRC*, and decreased (C) *FTH1* and (D) *FPN1* mRNA levels. Bars represent mean ± SEM with statistically significant differences determined by Student’s *t*-test (\**P* < 0.05, \*\**P* < 0.01, \*\*\*\**P* < 0.0001).

### Knockdown of *CYB561* mitigates the highly oncogenic phenotype of PC-3 cells

To ascertain if the effects of *CYB561* knockdown on NE phenotype and iron metabolism would translate to changes in the highly aggressive nature of PC-3 cells, we evaluated the consequences of *CYB561* knockdown on several hallmarks of cancer cell behavior. In proliferation assays, we observed a decrease in cell proliferation of PC-3 cells transduced with sh*CYB561* compared to scrambled shRNA control (Figs 7A-B). In a colony formation assay that is used as an indicator of the transforming ability of cells to survive very low cell densities and form colonies, we found that PC-3 sh*CYB561* cells formed smaller and fewer colonies relative to the scrambled shRNA control (Figs 7C-D). Lastly, in a wound-healing assay, PC-3 sh*CYB561* cells exhibited a slower migration rate demonstrated by the lower percentage of wound closure compared to the scrambled shRNA control (Figs 7E-F).

**Fig 7.**
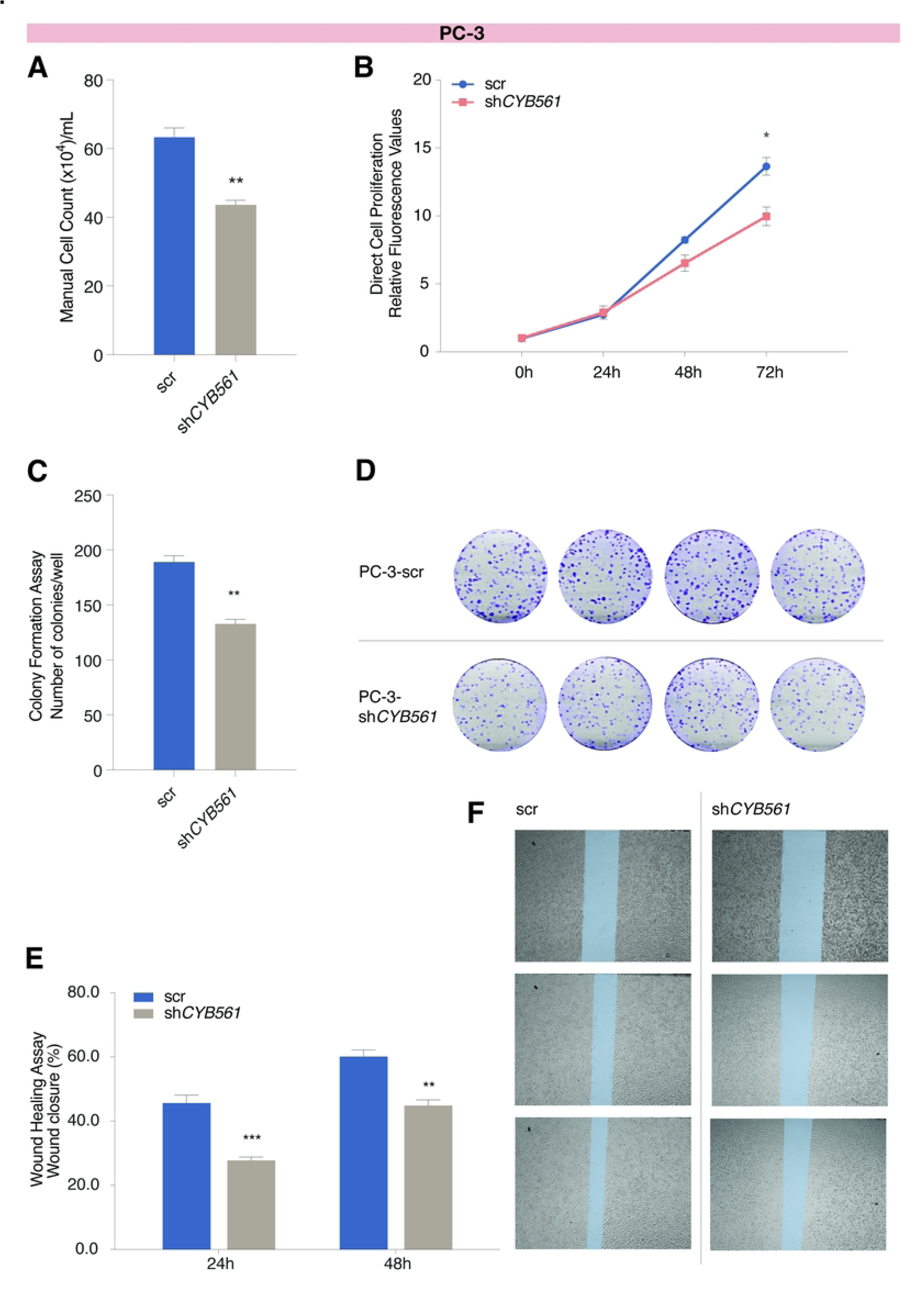
*CYB561* knockdown decreased the survival, proliferation, and migration rates of PC-3 cells. PC-3 cells transduced with sh*CYB561* had lower cell proliferation rates compared to the scrambled (scr) shRNA control in **(A)** trypan blue exclusion assay and **(B)** direct cell proliferation assay. Compared to the scr shRNA control, knocking down *CYB561* in PC-3 cells resulted in **(C-D)** fewer colonies in colony formation assays and **(E-F)** reduced the migration rate control as measured at 24 hr and 48 hr timepoints of a wound healing assay. Bars represent mean ± SEM with statistically significant differences determined by Student’s *t*-test (\**P* < 0.01, \*\**P* < 0.001, \*\*\**P* < 0.0001).

## Discussion

A growing body of evidence has implicated the role of ADT in promoting lineage plasticity during PCa progression and resistance development (36). The prevailing paradigm suggests that the resulting androgen-deficient environment from ADT exerts selective pressure on PCa cells, reducing their reliance on AR signaling and allowing the acquisition of NE-like features through transdifferentiation (37). This is supported by a focal increase in NE cell population adjacent to adenocarcinoma cells after hormone therapy (25), and the development of NE features and neuronal marker expression closely resembling t-NEPC when LNCaP cells are maintained in steroid-free conditions (22). Histologically, t-NEPC tumors present with small cell NE-like morphologies with neurite extensions, and express conventional NED markers such as SYP, ENO2, and CHGA (22, 38). Neuroendocrine cells influence the behavior of adjacent cells through a paracrine manner by producing a wide array of neuropeptides (39) which then promote the survival, proliferation, and migration of PCa cells (6). These studies implicate NE cells and neuropeptide signaling as having significant roles in therapy resistance and NEPC development.

Given the involvement of CYB561 in neuropeptide synthesis and its observed upregulation in CRPC and NEPC, we characterized *CYB561* gene expression pattern across different PCa tumor subtypes and PCa cell lines, explored the androgen-dependence of *CYB561* expression, demonstrated a role for CYB561 in neuropeptide synthesis and iron metabolism of CRPC, and more importantly, established the contribution of CYB561 to the transdifferentiation and maintenance of the highly aggressive NE phenotype in CRPC.

### CYB561 in metastatic prostate adenocarcinoma and castration resistance

Analysis of publicly available RNA-seq and copy number data revealed the highest level of *CYB561* expression in PCa tumors and increasing *CYB561* amplification frequencies with progression from CRPC to NEPC. In PCa cell lines, we also found higher CYB561 expression in metastatic PCa and NEPC models, in agreement with the specific enrichment of CYB561 in neuropeptide-containing tissues (40). The pattern of increasing *CYB561* amplification as the tumor progresses to CRPC and NEPC was also observed for *PAM* and genes encoding neuropeptide receptors further associating therapy resistance with the activation of neuropeptide signaling genes. These results are consistent with the known increase in PAM expression and activity in PC-3 cells, and lack of *PAM* expression and secreted neuropeptides in LNCaP cells indicating that the neuropeptide synthesis pathway is inactive in androgen-dependent LNCaP cells (41, 42).

Although *CYB561* is highly expressed in LNCaP cells relative to normal prostate cells, we found that *CYB561* expression is independent of androgen regulation. Nevertheless, knockdown of *CYB561* in LNCaP decreased androgen-dependent growth and enhanced sensitivity to Enz, suggesting that CYB561 contributes to the oncogenic potential of CRPC. We also observed *PAM* expression to be downregulated by androgen treatment and upregulated by acute and chronic CSS conditions in LNCaP cells. Notably, when we did the same treatment in 22Rv1 cells, a PCa cell line known to be less reliant on androgen for growth and representative of the progression from PRAD to NEPC, the androgen-dependent downregulation of *PAM* was abolished (17), consistent with our hypothesis that PAM may be negatively regulated by androgen signaling. In the future, it would be interesting to look at the individual contribution of PAM to NEPC progression. Along with our results that co-expression of *CYB561* and *PAM* is only detected in PC-3 cells, these findings provide evidentiary support that the neuropeptide activation function of *CYB561* may be more relevant in NEPC compared to androgen-dependent metastatic PCa.

While we cannot fully ascertain the relative levels of LIP without direct measurements of intracellular Fe^2+^ across cell lines, the pattern of IRG expression we observed points towards NEPC having higher LIP compared to PRAD. This is consistent with previous reports that showed metastatic PCa upregulating the autocrine hormone hepcidin to promote ferroportin degradation (43). This decrease in ferroportin expression effectively increases iron retention and promotes cancer cell survival (43). As for the significance of the ferrireductase function of CYB561 in PCa iron metabolism, knockdown of *CYB561* in LNCaP cells decreased Fe^2+^ concentration indicating that reduced CYB561 expression can lower cellular ferrireductase activity. These results, along with the sensitizing effect of CYB561 knockdown to Enz, may indicate a potential contribution of CYB561 ferrireductase function towards supporting tumor growth and therapy resistance through increased cellular iron load (9). Furthermore, these findings point to the benefit of using ferrireductase inhibitors in combination with established anti-PCa drugs to delay or prevent castration resistance.

### Functional role of CYB561 in the development and transdifferentiation towards NEPC

We induced NE transdifferentiation in LNCaP to determine changes in expression patterns of neuropeptide signaling and iron metabolism genes following NED. As expected, transdifferentiation increased expression of NED markers *SYP* (21), *ENO2* (22), and *CHGA* (22). However, *CYB561* expression did not change following NED. Similar results were observed when LNCaP cells were subjected to chronic steroid deprivation to model NEPC progression by long-term ADT, consistent with our observation that *CYB561* expression is not responsive to androgen manipulation. In contrast, *PAM* expression was upregulated by NED which is congruent with the reported increase in PAM expression as revealed by proteomics analysis (44). More importantly, the increase in PAM expression that occurs with NED supports the prevailing paradigm that androgens suppress signaling pathways that promote the NE phenotype (45).

To determine whether CYB561 is a key mediator of NED, we induced NE transdifferentiation in sh*CYB561*-transduced LNCaP cells. Knockdown of *CYB561* downregulated the expression of NED markers upon transdifferentiation, indicating a significant role for CYB561 in mediating lineage plasticity and facilitating NED in PCa. Interestingly, *CYB561* knockdown resulted in increased *PAM* transcript levels, possibly as a means to compensate for the reduced neuropeptide amidation resulting from lower ASC levels (46).

Knockdown of *CYB561* in LNCaP cells also dampened the transdifferentiation-induced increase in Fe^2+^ and total iron levels, demonstrating the significant contribution of CYB561 in the dysregulation of iron metabolism brought about by reprogramming towards NEPC. Relative to scrambled shRNA control cells, knockdown of *CYB561* led to a decrease in Fe^2+^ and total iron concentration that is supported by a decrease in *TFRC* and *FTH1* expression that may result in lower Fe^3+^ influx and storage, respectively, along with the increase in *FPN1* mRNA expression that may consequently increase Fe^2+^ efflux. Upon transdifferentiation, *CYB561* knockdown also lowered Fe^2+^ and total iron concentrations consistent with the observed IRG expression of pattern of lower *TFRC,* and higher *FTH1* and *FPN1* mRNA expression in *CYB561* knockdown cells relative to the scrambled shRNA control. These results emphasize the iron-seeking behavior of cancer cells as the disease progresses (9), and more importantly, the contribution of CYB561 ferrireductase activity towards t-NEPC.

While a definitive link between iron and NE transdifferentiation is lacking, the observation that Fe^2+^ is required for the activity of the chromatin modifying enzyme JARID1B provides a possible mechanism on how intracellular iron levels influence the chromatin landscape and consequently contribute to NEPC reprogramming (47). JARID1B, a histone H3 lysine 4 demethylase upregulated in several cancer types including PCa, requires Fe^2+^ and α-ketoglutarate as cofactors for its demethylase activity (47). One can then infer that changes in the availability and metabolism of these factors can affect histone demethylase activity and consequently the chromatin landscape.

### Functional role of CYB561 in the maintenance of the NEPC phenotype

In PC-3 cells, *CYB561* knockdown did not affect *PAM* expression but decreased the expression of NED markers *SYP*, *ENO2* and *CHGA*, suggesting a functional role for CYB561 in the maintenance of the NE phenotype. Notably, our paracrine signaling assay revealed that growth-promoting factors become limited when *CYB561* expression is reduced. While further analysis is needed to confirm that our results can be attributed to a decrease in the quantity of activated neuropeptides with *CYB561* knockdown, several studies have employed the use of conditioned media to investigate neuropeptide function (48, 49). Moreover, consistent with results of the paracrine signaling assay, we found that knocking down *CYB561* decreased the proliferation, survival, and migration of PC-3 cells. Taken together, these results provide further evidence that upregulation of genes relevant to neuropeptide synthesis, such as *PAM* and *CYB561*, is an important mechanism employed by PCa to circumvent ADT, develop castration resistance, and induce and maintain the NE phenotype.

The comparative pattern of IRG expression across the different PCa cell lines are consistent with a consequent increase of LIP levels in PC-3 in comparison to LNCaP and 22Rv1 cells. We were able to demonstrate that similar to LNCaP cells, knocking down *CYB561* in PC-3 cells can effectively reduce Fe^2+^ concentrations and that PC-3 cells may compensate by upregulating the influx of iron via an in increase *TFRC* expression, and downregulating iron storage and efflux by decreasing *FTH1* and *FPN1*, respectively (9). Our results show and validate, for the first time, that apart from its well-known function in ASC recycling and neuropeptide synthesis, CYB561 also affects the iron metabolism of PCa cells.

## Conclusion

In summary, our findings provide initial evidence that through the coordinate involvement of CYB561 in neuropeptide signaling and iron metabolism, these seemingly unrelated pathways can cooperatively contribute towards PCa progression. We propose that the ferrireductase function of CYB561 initially supports the metastatic and proliferation potential of PRAD and CRPC by increasing iron load and available LIP. Upon hormone castration by ADT, the induced expression of PAM enables CYB561 to contribute to the activation of the neuropeptide amidation pathway through its role in recycling ASC. By integrating these two pathways, CYB561 sustains the NE phenotype and promotes a more aggressive cancer cell behavior by supporting cell survival, proliferation, and migration *in vitro* (Fig 8). The complex effects triggered by mere manipulation of *CYB561* expression level in PCa cell models suggest that CYB561 has a significant contribution in facilitating PCa tumor behavior. Furthermore, our results significantly expand the current knowledge on iron dysregulation and PCa, providing a link between androgen deprivation, intracellular iron levels and NEPC progression. To our knowledge, this is the first study to demonstrate the dual role of CYB561 in the context of PCa cell behavior and to provide a functional basis for its observed upregulation in the advanced stages of the disease.

**Fig 8.**
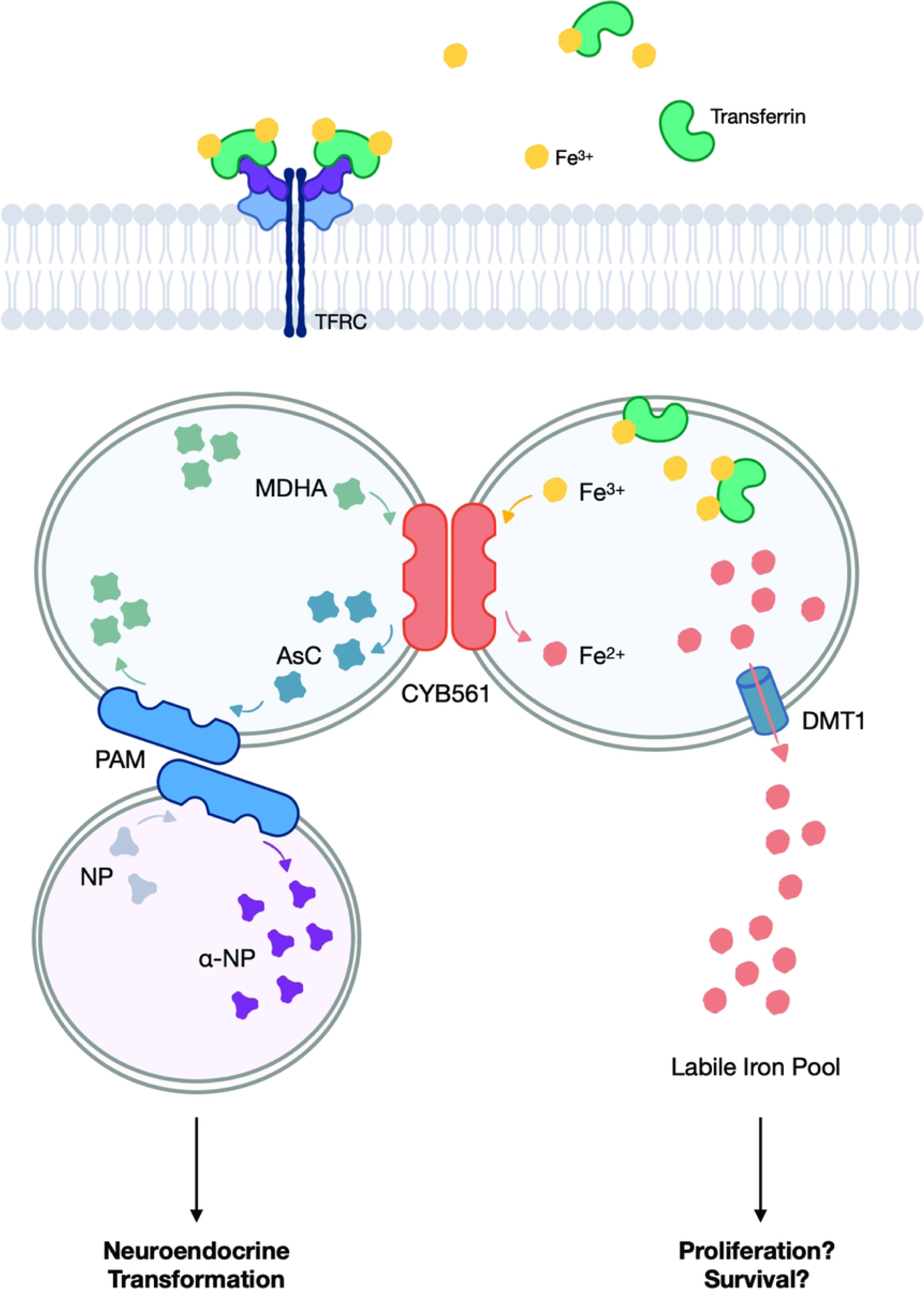
*CYB561* promotes and sustains the more aggressive NEPC phenotype through its role in ascorbate recycling and iron homeostasis. CYB561 replenishes the ascorbic acid pool which is then used by PAM to activate neuropeptides that promote transformation towards NEPC. Additionally, CYB561 increases the availability of intracellular iron which is essential for regulating cellular activities related to proliferation and survival.

## Supporting Information

**S1 Fig. Chronic steroid starvation of LNCaP cells induces NE transdifferentiated phenotype. (A)** LNCaP cells were maintained in complete growth media (FBS) and charcoal steroid-stripped (CSS) media for 10 mo (chronic CSS; cCSS). Chronic steroid starvation increased *SYP* mRNA expression. **(B)** Representative images (200X magnification) of LNCaP-FBS and LNCaP-cCSS at 90 days of cell culture maintenance are shown. The LNCaP-cCSS cells developed long cytoplasmic extensions resembling a neuron-like morphology. Bars represent mean ± SEM with statistical significance indicated by asterisks in Student’s *t*-test (\**P* < 0.05).

**S2 Fig. *In silico* gene expression analyses of selected neuropeptide receptor genes in PCa.** Gene amplification frequency of the neuropeptide receptor genes: **(A)** gastrin releasing peptide receptor (*GRPR*), **(B)** calcitonin receptor like receptor (*CALCRL*), **(C)** neurotensin receptor 1 (*NTSR1*), **(D)** neuropeptide Y receptor (*NPY1R*), **(E)** vasopressin receptor 1A (*AVPR1A*) were examined in different types of PCa using publicly available RNA-seq and microarray copy number data (25), and visualized using the online visualization tool cBioPortal (27, 34). All five genes examined have higher amplification frequencies in the more advanced CRPC and NEPC stages relative to early-stage PRAD.

**S3 Fig. Basal expression of *CYB561* homologs in normal and PCa cell lines. (A)** The duodenal *CYB561* homolog, *DCYTB*, shows high expression levels in the normal cell lines RWPE-1 and PNT1A. **(B)** The expression of lysosomal homolog, *LCYTB*, is similar across all cell lines analyzed. **(C)** The annotated tumor suppressor *101F6* exhibits high expression in LNCaP cells. Bars represent mean ± SEM with statistically significant differences determined through one-way ANOVA followed by Tukey’s post-hoc test (*P* < 0.05; means with the same letter are not statistically different).

**S4 Fig. Analysis of androgen-dependent expression of *CYB561* and *PAM*. (A)** DHT treatment of LNCaP cells grown for 48 hr in complete growth media (FBS) or acute charcoal steroid-stripped (aCSS) media induced *KLK3* mRNA expression which was abolished in the presence of Enzalutamide (+Enz) (one-way ANOVA; FBS: *P* < 0.0001; aCSS: *P* < 0.0001; Student’s *t*-test; Veh: *P* < 0.0001; +Enz: *P* = 0.0003). **(B)** *CYB561* mRNA expression was not affected by hormone treatment in both media setups (one-way ANOVA; FBS: *P* = 0.1483; aCSS *P* = 0.1546). **(C)** *PAM* mRNA levels increased with aCSS (Student’s *t*-test; Veh: *P* < 0.0001; DHT: *P* = 0.003; +Enz: *P* = 0.0155) relative to the cells grown in FBS. Treatment with DHT dampened the increase in *PAM* expression and this was reversed by Enz treatment (one-way ANOVA; aCSS: *P* = 0.0281). **(D-F)** *KLK3, CYB561,* and *PAM* mRNA expression were measured in LNCaP cells grown and maintained in complete growth media (FBS) or chronic charcoal steroid-stripped (cCSS) media for 10 mo. Maintenance in cCSS depleted the expression of **(D)** *KLK3* mRNA, **(E)** did not affect *CYB561* mRNA expression, and **(F)** increased *PAM* mRNA levels. Bars represent mean ± SEM with statistically significant differences determined through one-way ANOVA for effect of hormone treatment within the same growth media followed by Tukey’s post-hoc test (*P* < 0.05; means with the same letter are not statistically different) and Student’s *t-*test for the individual effects of growth media between the same hormone treatment (^#^*P* < 0.05, ^##^*P* < 0.01, ^###^*P* < 0.001, and ^####^*P* < 0.000), or for the effect of growth media (\**P* < 0.0001).

**S5 Fig. Validation of CYB561 knockdown in LNCaP cells. (A)** Effective CYB561 knockdown was achieved at the **(A)** mRNA level by 74.24% as determined by RT-qPCR that was reflected at **(B)** the protein level as determined by western blot analysis in LNCaP cells transduced with sh*CYB561* compared to the scrambled (scr) shRNA control. **(C)** Hormone treatment did not alter *CYB561* expression in scr shRNA and sh*CYB561* cells. Bars represent mean ± SEM with statistically significant differences determined through Student’s *t-*test for the effect of *CYB561* knockdown (\**P* < 0.0001) or the effect of *CYB561* knockdown between the same hormone treatment (^#^*P* < 0.0001).

**S6 Fig. Depletion of CYB561 lowered the sensitivity of LNCaP to androgen-stimulated and Enz-mediated effects on cell proliferation. (A-C)** Proliferation rates of transduced LNCaP cells upon hormone treatment were measured every 72 hr over the course of one week. Knockdown of *CYB561* reduced the proliferative effects of DHT at **(A)** 72 hr and **(B)** 120 hr post-hormone treatment. Interestingly, *CYB561* knockdown enhanced the repressive effect of Enz on cell proliferation at the **(C)** 168 hr timepoint. Data points and bars represent mean ± SEM with statistically significant differences determined through one-way ANOVA followed by Tukey’s post-hoc test for effect of hormone treatment (*P* < 0.05; means with the same letter are not statistically different) and Student’s *t-*test for the effect of *CYB561* knockdown between the same hormone treatment (^#^*P* < 0.01)

**S7 Fig. *CHGA* mRNA expression decreases with *CYB561* knockdown.**

LNCaP cells transduced with the scrambled (scr) shRNA control and sh*CYB561* were grown and maintained in complete media (control) or transdifferentiation media for 14 days. Transdifferentiation induced a slight increase in relative *CHGA* mRNA levels in scr control cells (two-way ANOVA; Treatment factor: *P* = 0.0315; Knockdown factor: *P* = 0.0714). Bars represent fold-change values ± SEM of ΔCt with statistically significant differences determined through two-way ANOVA for main effects of treatment and *CYB561* knockdown and Student’s *t-*test for the individual effects of treatment within an shRNA type (\**P* < 0.01).

**S8 Fig. Basal expression of iron-responsive genes (IRGs) in prostate cancer cell lines and effect of *CYB561* knockdown and transdifferentiation on total iron concentration and IRG expression in LNCaP cells.** (A) *TFRC* mRNA is highly expressed in LNCaP, 22rv1, and PC-3 cancer cell lines compared to normal prostate epithelial models RWPE-1. (B) *FTH1* mRNA level is highest in PC-3 cells while (C) *FPN1* mRNA is highly expressed in PNT1A and 22rv1 cells. **(D)** LNCaP cells transduced with the scrambled (scr) shRNA control and sh*CYB561* were grown and maintained in complete media (control) or transdifferentiation media for 14 days followed by quantitation of total iron concentration and IRG expression. The total iron concentration of LNCaP cells increased with transdifferentiation and this effect was dampened with *CYB561* knockdown (two-way ANOVA; Treatment factor: *P* < 0.0001; Knockdown factor: *P* < 0.0001). **(E-F)** Expression of IRGs was assessed in LNCaP maintained in complete growth media (FBS) control or chronic charcoal steroid-starved (cCSS). Chronic steroid starvation **(E)** did not affect *TFRC* mRNA levels but **(F)** led to an increase in *FTH1* mRNA expression. **(G-H)** LNCaP cells were treated with vehicle (Veh), 10 nM DHT, or 10 nM DHT plus 10 uM Enz for 24 hr before harvest for RT-qPCR analysis. (G) *TFRC* and (H) *FTH1* mRNA expression remained unchanged across all treatments. (I) *STEAP2* mRNA levels were measured in the transdifferentiated LNCaP cells by RT-qPCR. Transdifferentiation resulted in an increase in *STEAP2* expression which was not altered by *CYB561* knockdown (two-way ANOVA; Treatment factor: *P* < 0.0001; Knockdown factor: *P* = 0.2520). Bars represent mean ± SEM with statistically significant differences determined through one-way ANOVA followed by Tukey’s post-hoc test (*P* < 0.05; means with the same letter are not statistically different), or two-way ANOVA for main effects of media treatment and *CYB561* knockdown, and Student’s *t-*test for the individual effects of media treatment (\**P* < 0.001, \*\**P* < 0.0001) and *CYB561* knockdown between the same treatment (^#^*P* < 0.01).

**S9 Fig. Gene expression analysis and total iron concentration in *CYB561* knockdown PC-3 cells. (A-B)** Validation of *CYB561* knockdown in PC-3 cells showed that **(A)** *CYB561* mRNA expression was reduced by 74.23% at the mRNA level which was **(B)** reflected at the level of protein expression in PC-3 cells transduced with sh*CYB561* compared to the scrambled (scr) shRNA control. Knockdown of *CYB561* **(C)** decreased *CHGA* mRNA expression but **(D)** did not affect total iron concentration in PC-3 cells. Bars represent mean ± SEM for *CYB561* mRNA and total iron concentration plots while bars represent relative fold-change values ± SEM of the computed ΔCt values for *CHGA* plots. Statistically significant differences due to *CYB561* knockdown were determined through Student’s *t-*test (\**P* < 0.0001)

**Table S1.**
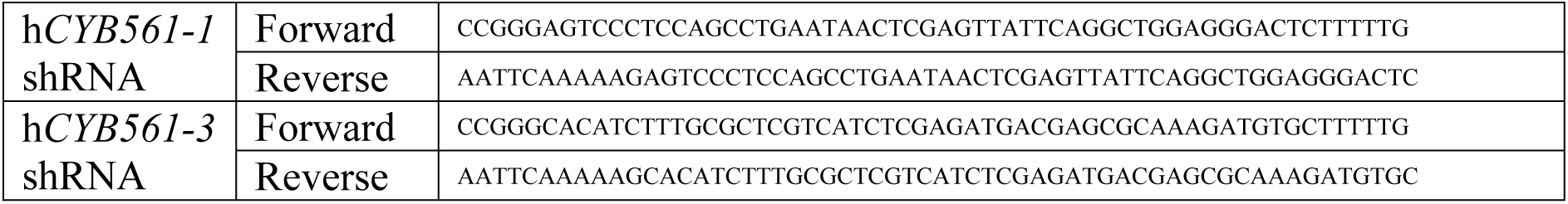
Primers used to generate the pLKO.1-sh*CYB561* construct for lentiviral transduction.

## Acknowledgement

We thank Dr. Diane Robins (University of Michigan Medical School) for providing the RWPE-1 and PC-3 cell lines, and for her thorough review of our manuscript. We also thank the Philippine Council for Health Research and Development of the Department of Science and Technology for establishing the Genome Editing Facility that allowed us to conduct lentiviral work.

